# HB-EGF and EGF infusion following CNS demyelination mitigates age-related decline in regeneration of oligodendrocytes from neural precursor cells originating in the ventricular-subventricular zone

**DOI:** 10.1101/2024.02.26.582092

**Authors:** Kaveh Moradi, Stanislaw Mitew, Yao Lulu Xing, Tobias D. Merson

## Abstract

In multiple sclerosis (MS), chronic demyelination initiated by immune-mediated destruction of myelin, leads to axonal damage and neuronal cell death, resulting in a progressive decline in neurological function. The development of interventions that potentiate remyelination could hold promise as a novel treatment strategy for MS. To this end, our group has demonstrated that neural precursor cells (NPCs) residing in the ventricular-subventricular zone (V-SVZ) of the adult mouse brain contribute significantly to remyelination in response to central nervous system (CNS) demyelination and can regenerate myelin of normal thickness. However, aging takes its toll on the regenerative potential of NPCs and reduces their contribution to remyelination. In this study, we investigated how aging influences the contribution of NPCs to oligodendrogenesis during the remyelination process and whether the delivery of growth factors into the brains of aged mice could potentiate the oligodendrogenic potential of NPCs. To enable us to map the fate of NPCs in response to demyelination induced at different postnatal ages, *Nestin-CreER^T2^*; *Rosa26-LSL-eYFP* mice were gavaged with tamoxifen at either 8 weeks, 30 weeks or one year of age before being challenged with cuprizone for a period of six weeks. Using osmotic minipumps, we infused heparin-binding EGF-like growth factor (HB-EGF), and/or epidermal growth factor (EGF) into the cisterna magna for a period of two weeks beginning at the peak of cuprizone-induced demyelination (n=6-8 mice per group). Control mice received artificial cerebrospinal fluid (vehicle) alone. Mice were perfused six weeks after cuprizone withdrawal and the contribution of NPCs to oligodendrocyte regeneration in the corpus callosum was assessed. Our data reveal that although NPC-derived oligodendrocyte generation declined dramatically with age, this decline was partially reversed by growth factor infusion. Notably, co-infusion of EGF and HB-EGF increased oligodendrocyte regeneration twofold in some regions of the corpus callosum. Our results shed light on the beneficial effects of EGF and HB-EGF for increasing the contribution of NPCs to remyelination and indicate their therapeutic potential to combat the negative effects of aging upon remyelination efficacy.

## Introduction

Multiple sclerosis (MS) is characterized by immune-mediated destruction of myelin, resulting in axonal damage and neuronal cell death, which leads to a progressive decline in neurological function ^1^. In response to myelin loss, a process known as remyelination is initiated, which restores myelin sheaths to demyelinated axons through the generation of new myelin-forming oligodendrocytes ^2^. New oligodendrocytes are generated via the differentiation of parenchymal oligodendrocyte progenitor cells (pOPCs) or neural precursor cells (NPCs) born in the ventricular-subventricular zone (V-SVZ) ^3–5^.

Whilst remyelination appears to be efficient in the early stages of MS, this regenerative process declines with disease chronicity ^6, 7^. Aging is one of the most important risk factors for remyelination failure ^8, 9^. Whilst the effect of aging on pOPCs has been explored ^10–13^, it is not clear how aging may affect the contribution of NPCs to remyelination. Although some studies suggested that proliferation of NPCs declines with aging ^14–16^, other studies have shown that the proliferation of these cells does not significantly change over time ^17, 18^. Moreover, these studies concentrated solely on age-related alterations in the neurogenic potential of NPCs and did not examine any associated changes in the oligodendrogenic capacity of NPCs during remyelination.

The microenvironment of the V-SVZ regulates the fate of NPCs. Growing data suggests that the aged V- V-SVZ niche is a major factor in driving NPCs in the quiescence stage, leading to a decrease in the neurogenic potential of these cells in the aged brain ^19, 20^. One characteristic of the aged V-SVZ niche is a change in the expression level of growth factors and their receptor such as transforming growth factor-α (TGF-α), epidermal growth factor (EGF), epidermal growth factor receptor (EGFR), insulin-like growth factor-1 (IGF-1) and vascular endothelial growth factor (VEGF) ^21–25^. Reduced EGF signaling is aged V-SVZ associated with decreased NPC-derived neurogenesis ^21^. Some studies suggested that EGF or heparin-binding EGF-like growth factor (HB-EGF) infusion improves the neurogenic potential of NPCs in aged mice ^22, 26^. In contrast, others showed that EGF-responsive cells in the V-SVZ are transient amplifying cells (Type C cells) and EGF administration to these cells arrests their neurogenic capacity, converting them into multipotent stem cells from neurogenic precursors ^27^. The level of EGF in the cerebrospinal fluid (CSF) of MS patients is lower than in healthy people, which might be associated with remyelination failure in these patients ^28^. Several studies have suggested that EGF or HB-EGF can induce NPCs to produce more oligodendrocytes, which can improve remyelination after demyelination insults ^29, 30^. Further investigation is required to determine whether these growth factors induce NPCs to differentiate into oligodendrocytes^30^, or into other cell lineages ^22, 29^. In addition, none of these studies used old animal models to see the reaction of old NPCs to these growth factors.

In this study, we examined the contribution of NPCs to remyelination using *Nestin-CreER^T2^:Rosa26-eYFP* mice at 8 weeks, 30 weeks, and 1 year of age to assess the effects of aging on their oligodendrogenic potential. In addition, we examined how the administration of EGF and/or HB-EGF into the CSF of mice affects the behavior of NPCs following a demyelinating insult. Additionally, we investigated whether these growth factors can enhance the oligodendrogenic potential of NPCs following cuprizone challenge.

## Materials and Methods

### Animals

All animal experiments were conducted following the National Health and Medical Research Council guidelines and were approved by the Monash Animal Research Platform (MARP) Animal Ethics Committee (ethics number 172). Experimental cohorts were generated by crossing *Nestin-CreER^T2^ Line 5-1* transgenic mice ^31^ and *Rosa26-eYFP* reporter mice ^32^, both maintained on a *C57BL/6J* background, to yield *Nestin-CreERT2 line 5-1: Rosa26-eYFP* double-transgenic mice (denoted *Nestin:YFP*).

### Tamoxifen administration

Nestin^+^ NPCs were genetically labeled using *Nestin:YFP* mice. To achieve this, the mice were given tamoxifen (Sigma Cat# T5648) via oral gavage at a dose of 0.3 g/kg/d for four consecutive days. The tamoxifen was prepared at 40 mg/ml in corn oil (Sigma Cat# C8267). For the vehicle control group, corn oil without tamoxifen was administered.

### Induction of demyelination

One week after the last tamoxifen gavage, *Nestin:YFP* mice that were 8 weeks, 30 weeks, or 1 year old were fed with standard rodent chow mixed with 0.2% cuprizone (w/w; bis-cyclohexanoneoxaldihydrazone; Sigma Cat# C9012) to induce demyelination. The mice used for EGF and/or HB-EGF infusion experiments (n=6/group) were on the cuprizone diet for 6 consecutive weeks and then returned to a normal diet for 6 weeks of recovery. The mice used to investigate the extent of demyelination (n=4-5/group) were on the cuprizone diet for 5 consecutive weeks and were then sacrificed. For the latter experiment, the controls remained on a normal diet throughout the experiment.

### EGF and/or HB-EGF infusion into the CSF

Following a 5-week cuprizone challenge, the *Nestin:YFP* mice were given isoflurane anesthesia, and osmotic mini-pumps (ALZET® Model 2002) were implanted subcutaneously. These mini-pumps were used to infuse growth factors directly into the cerebrospinal fluid (CSF) through the cisterna magna at a flow rate of 0.25 µl/hour. The osmotic mini-pumps contained 100 µl of either EGF (66.7 µg/mL, Millipore Cat# 01-102), HB-EGF (20 µg/mL, R and D systems Cat# 259-HE-050/CF), EGF+HB-EGF (66.7 µg/mL and 20 µg/mL, respectively), or aCSF (vehicle control) (TOCRIS Bioscience Cat# 3528). This equates to 400 ng/day EGF, a dose previously demonstrated to induce oligodendrogenesis of neural stem cells in the V-SVZ ^30^, and 120 ng/day HB-EGF, a dose previously demonstrated to restore neurogenesis in the V-SVZ in the aged brain ^22^. The mini-pumps were removed after 2 weeks of implantation.

### EdU administration

To study the impact of EGF and HB-EGF infusion on rapidly dividing neural stem cells, we conducted experiments on 8-week-old *Nestin:YFP* mice treated with cuprizone. Each group (n=5-6) of mice received a single injection of EdU (50 mg/kg) at a dilution of 5 mg/ml in saline, 3 days after implanting the osmotic mini-pumps. The injection was given at a rate of 10 μl/g body weight, i.p. The mice were humanely sacrificed 2 hours after the EdU injection.

### Perfusion/fixation

Mice were anesthetized with sodium pentobarbitone (100 mg/kg, i.p.) and perfused with 4% paraformaldehyde in 0.1 M PBS (pH 7.4). The brains were collected and post-fixed in 4% PFA/PBS at 4°C for 2 hours then cryoprotected for 24 hours in 20% sucrose/PBS at 4°C. Brain tissue was then cut into coronal sections with a thickness of 10 microns and stored at −80°C until processed for immunohistochemistry.

### Immunohistochemistry

IHC was performed as previously described ^4^. Primary antibodies were as follows: mouse anti-APC (CC1) (1:100, Cabiochem Cat# D35078, RRID:AB_2057371), Guinea-pig anti-DCX (1:500, Chemicon Cat# AB5910, RRID:AB_2230227), mouse anti-GFAP (1:400, Chemicon Cat# MAB360, RRID:AB_11212597), mouse anti-MBP (1:1000, Millipore Cat# NE1018, RRID:AB_2140494), chicken anti-PLP (1:500, Aves Lab Cat# PLP, RRID:AB_2313560), chicken anti-GFP (1:1000, Aves Lab Cat# GFP-1020, RRID:AB_10000240), rabbit anti-SOX10 (1:500, Chemicon Cat# AB5727, RRID:AB_2195375), goat anti-Pdgfrα (1:50, R and D Systems Cat# AF1062, RRID:AB_2236897), rabbit anti-ALDH1L1 (1:1000, Abcam Cat# ab7117, RRID:AB_954955), mouse anti-PCNA (1:500, Abcam Cat# ab29, RRID:AB_303394), rabbit anti-OLIG2 (1:100, IBL-America Cat# JP18953, RRID:AB_1630817) and rabbit anti-MCM2 (1:200, Abcam Cat# ab4461, RRID:AB_304470). Staining was revealed using appropriate secondary antibodies raised in donkey including anti-chicken conjugated to FITC (1:200, Jackson ImmunoResearch Labs Cat# 703-095-155, RRID:AB_2340356), anti-chicken conjugated to Alexa Flour 488 (1:200, Jackson ImmunoResearch Labs Cat# 703-545-155, RRID:AB_2340375), anti-rabbit conjugated to Alexa Flour 647 (1:200, Jackson ImmunoResearch Labs Cat# 711-605-152, RRID:AB_2492288), anti-rabbit conjugated to TRITC (1:200, Jackson ImmunoResearch Labs Cat# 711-025-152, RRID:AB_2340588), anti-rabbit conjugated to FITC (1:200, Jackson ImmunoResearch Labs Cat# 711-095-152, RRID:AB_2315776), anti-mouse conjugated to TRITC (1:200, Jackson ImmunoResearch Labs Cat# 715-025-150, RRID:AB_2340766), anti-mouse conjugated to Alexa Flour 647 (1:200, Jackson ImmunoResearch Labs Cat# 705-606-151, RRID:AB_2340866), anti-mouse conjugated to Alexa Flour 594 (1:200, Thermo Fisher Scientific Cat# A21203, RRID:AB_141633), anti-goat conjugated to Alexa Flour 594 (1:200, Jackson ImmunoResearch Labs Cat# 705-156-147, RRID:AB_141788), anti-goat conjugated to Alexa Flour 555 (1:200, Thermo Fisher Scientific Cat# A21432, RRID:AB_141788) and anti-guinea pig conjugated to Biotin-SP (1:200, Jackson ImmunoResearch Labs Cat# 706-065-148, RRID:AB_2340451). Sections incubated with biotinylated secondary antibody were rinsed and further incubated with the tertiary antibody, streptavidin-Brilliant Violet 480 (1:100, BD Biosciences Cat# 564876, RRID:AB_2869619). Cell nuclei were stained using Hoechst 33342 (0.1 µg/ml, Invitrogen Cat# H1399). To evaluate the integration of EdU in neural stem cells, the brain sections were initially subjected to immunostaining with GFAP and MCM2 antibodies. Afterward, the EdU detection was performed using the Alexa Fluor-647 Click-iT EdU Cell Proliferation Assay kit (Invitrogen Cat# C10424), following the manufacturer’s protocol.

### Image acquisition and quantifications

Laser scanning confocal microscopy (Zeiss LMS 780, Germany) equipped with a digital camera was used to capture micrographs required for quantifications. For analyzing the regional distribution of cells in the entire CC, tile scanning was performed using up to four different fluorophores with minimal overlap, for immunostaining. To detect more than four fluorophores or to distinguish overlapping fluorophores, the emission fingerprinting technique was applied. Spectral reflectance imaging was conducted on non-stained sections to visualize myelin in the CC. These images were then imported to ImageJ software (NIH) to quantify different cell populations, fluorescent intensity of myelin-specific proteins, or the total amount of myelin present in the CC. Nine sections were examined for each animal at each of the rostral, middle, and caudal segments of the CC, between levels 1.10 and 1.94 mm relative to Bregma. In all three segments, we quantified cell density in the mediolateral axis of the corpus callosum relative to the distance from the dorsolateral corner of the V-SVZ. To determine the amount of myelin-specific proteins or the total amount of myelin, we measured the mean gray value using ImageJ software (NIH) and reported it as optical density (OD). The quantitative data is presented as mean ± SEM. All cell counts and measurements were performed in a blinded manner.

### Experimental design and statistical analysis

Mice were selected randomly for each experiment. The gender and number of animals used for each group in experiments is provided in the corresponding figure legend. All data in the histograms are presented as mean ± SEM. Statistical analysis was performed using GraphPad Prism software 10.0 (GraphPad, RRID:SCR_002798). Differences between two groups were evaluated using the Student’s unpaired t-test. Meanwhile, differences between multiple groups were assessed using either a one-way or two-way ANOVA followed by Sidak’s or Tukey’s multiple comparisons tests. A P-value less than 0.05 was considered statistically significant. No animal was excluded from the analysis.

## Results

### Tamoxifen-induced recombination in the V-SVZ of Nestin:YFP mice enables NPC fate mapping across the lifespan

We have previously demonstrated that NPCs make a significant contribution to oligodendrocyte regeneration in the corpus callosum of mice following cuprizone-induced demyelination ^4^. In that study, we used young adult *Nestin:YFP* mice, aged 8 weeks, to track the fate of NPCs after cuprizone challenge. In the present study, we aimed to investigate whether the ability of NPCs to regenerate oligodendrocytes in the corpus callosum changes when demyelination is induced at different postnatal ages. Specifically, we sought to compare the oligodendrogenic potential of NPCs following demyelination induced at 8 weeks, 30 weeks, or 1 year of age.

To assess whether *Nestin:YFP* mice can be used for consistent fate-mapping of NPCs regardless of postnatal age, we first evaluated the efficiency of Cre recombination among dividing NPCs localized within the V-SVZ. We administered tamoxifen to mice at 8 weeks, 30 weeks, or 1 year of age and sacrificed them one week later. We determined the efficiency of Cre recombination by calculating the percentage of mitotic V-SVZ cells (PCNA^+^) that co-expressed YFP across five regions of the V-SVZ (**Figure 1A-C**). PCNA^+^ cells were observed in the dorsolateral, lateral, ventral, and medial V-SVZ (**Figure 1D**). No PCNA^+^ cells were detected in the dorsal wall of the lateral ventricles in any mice. Our analysis revealed no significant effect of postnatal age on recombination efficiencies among PCNA^+^ cells (*F* (2, 59) = 0.7523, *p*=0.4758). The mean recombination efficiency among PCNA^+^ cells was 77.8 ± 2.5%, 78.8 ± 3.1%, and 74.4 ± 3.2% at 8 weeks, 30 weeks, and 1 year, respectively. These data suggest that age does not significantly affect Cre recombination efficiency in *Nestin:YFP* mice. Thus, we concluded that the transgenic line was suitable for comparing the fate of NPCs across different postnatal ages.

**Figure 1.**
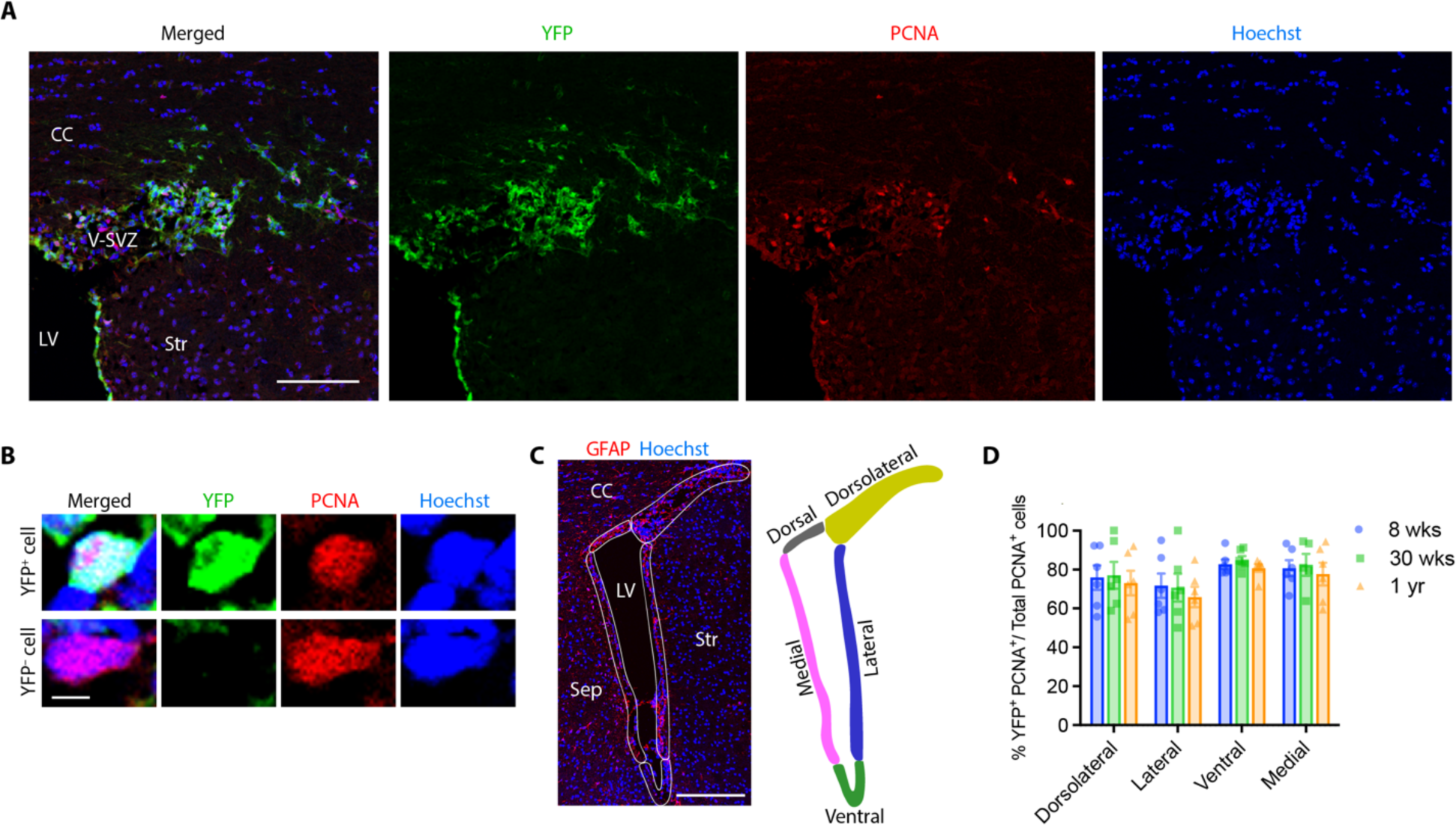
The efficiency of Cre-mediated recombination among dividing NPCs within subdomains of the V-SVZ of *Nestin:YFP* mice was similar across postnatal age. **A**, Confocal micrograph of the dorsolateral corner of the lateral ventricle of a 30-week-old *Nestin:YFP* mouse assessed 1 week after tamoxifen administration reveals abundant YFP^+^ PCNA^+^ cells (recombined mitotic NPCs) and a small number of YFP^−^ PCNA^+^ cells (non-recombined mitotic NPCs). **B**, Examples of YFP^+^ PCNA^+^ and YFP^−^ PCNA^+^ cells in the V-SVZ of a tamoxifen-gavaged *Nestin:YFP* mouse observed at high magnification. **C**, Coronal micrograph of the V-SVZ of the rostral forebrain. Demarcated areas show the five subdomains in which Cre recombination efficacy was assessed, namely the dorsolateral corner, lateral wall, ventral wall, medial wall, and dorsal wall of the lateral ventricles. The section was immunolabeled against GFAP and counterstained with Hoechst. **D**, Efficiency of Cre-mediated recombination among NPCs in the four regions of the V-SVZ where PCNA^+^ cells observed was estimated by determining the %YFP^+^ PCNA^+^ cells among total PCNA^+^ cells within each region. Data were analyzed by two-way ANOVA with Bonferroni’s post hoc test, n=6 mice/group. Data represent mean ± SEM. Scale bars: 100 µm (A), 5 µm (B), 200 µm (C). CC, corpus callosum; LV, lateral ventricle; Sep, septum; Str, striatum; V-SVZ, ventricular-subventricular zone.

### Contribution of NPCs to oligodendrogenesis following cuprizone challenge declined sharply with age

To investigate the effect of postnatal age on the generation of NPC-derived oligodendrocytes after demyelination, *Nestin:YFP* mice aged 8 weeks, 30 weeks, or 1 year were administered tamoxifen, then fed cuprizone for 6 weeks to induce demyelination and allowed to recover after cuprizone withdrawal for a further 6 weeks before perfusion/fixation (**Figure 2A**). Coronal sections including the rostral, middle and caudal segments of the corpus callosum were collected and processed immunohistochemically to detect YFP, SOX10 (a pan-oligodendroglial lineage marker), CC1 (an oligodendrocyte marker) and Hoechst (**Figure 2B,C**). Quantitation of total YFP^+^ cell density in the corpus callosum at rostral, middle and caudal segments revealed a marked age-associated decline in the density of NPC-fate mapped cells in the corpus callosum (**Figure 2D**). A two-way ANOVA was performed to analyze the effects of age and rostrocaudal position on YFP^+^ cell density. There was a statistically significant interaction between the effects of age and rostrocaudal position (*F* (4, 35) = 69.68, *p*<0.0001). Simple main effects analysis showed that both age and rostrocaudal position had a statistically significant effect on YFP^+^ cell density (*p*<0.0001).

**Figure 2.**
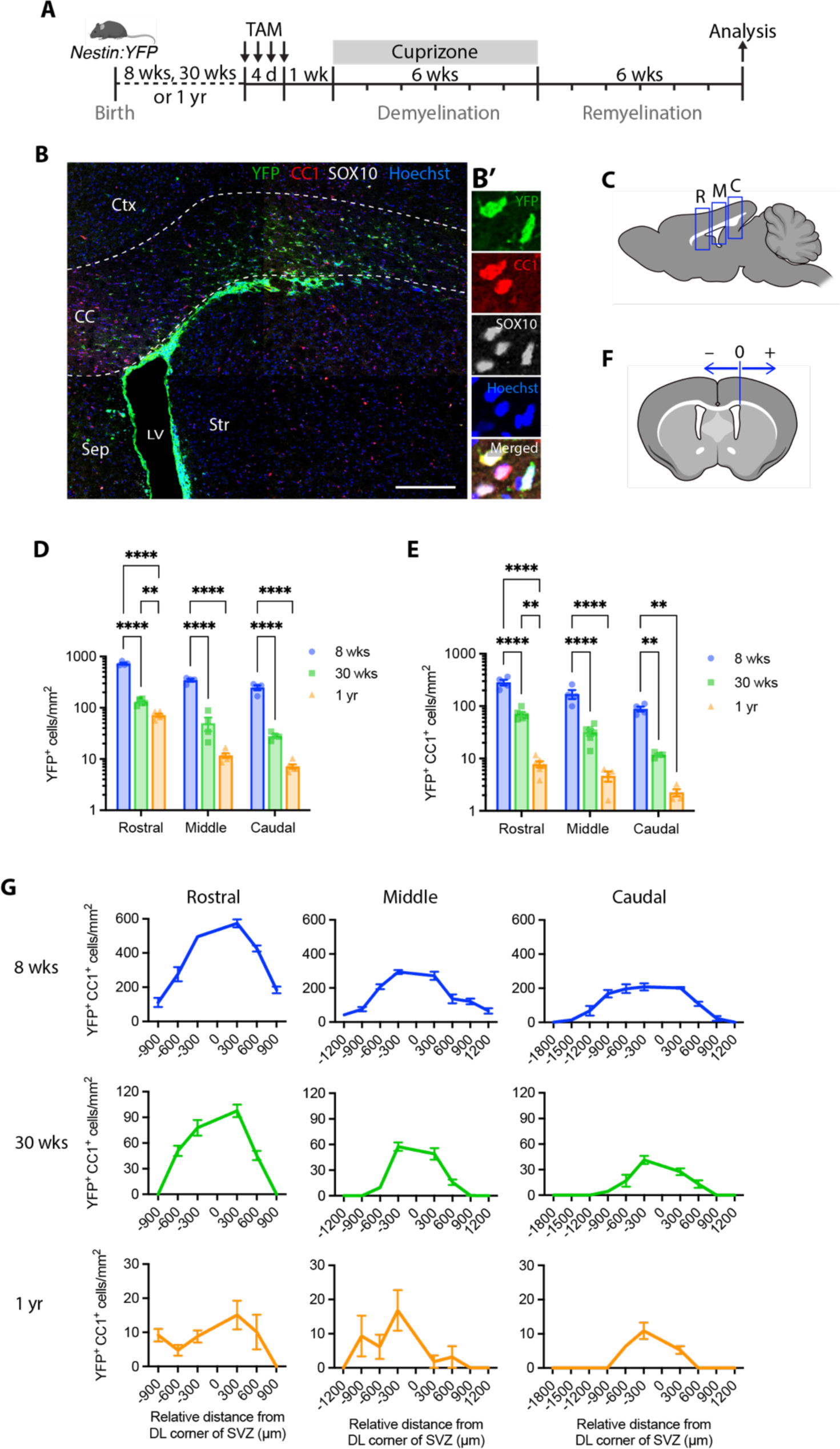
Aging dramatically reduced the extent to which NPCs contributed to oligodendrogenesis in the corpus callosum following cuprizone-induced demyelination. **A**, Experimental timeline for *Nestin:YFP* mice indicating time points for tamoxifen gavage, cuprizone challenge and tissue collection. **B**, Confocal micrograph of a coronal brain section from a *Nestin:YFP* mouse administered tamoxifen at 30 weeks of age then fed cuprizone for 6 weeks and perfused at 6 weeks recovery post cuprizone withdrawal. Immunohistochemistry was performed to detect YFP (green), CC1 (red) and SOX10 (grey). Sections were counterstained with Hoechst. The dashed line delineates the region of the corpus callosum. **B’**, Representative high magnification micrograph of labeled cells in the corpus callosum. **C**, Schematic representation of a sagittal section of the adult mouse brain illustrating the rostral (R), middle (M), and caudal (C) segments of the corpus callosum that were examined (relates to plots in D and E). **D**,**E**, Histograms showing the mean density of total YFP^+^ cells (NPC fate-mapped cells, D) or YFP^+^ CC1^+^ cells (NPC-derived oligodendrocytes, E) detected in rostral, middle, and caudal segments of the corpus callosum of cuprizone-challenged *Nestin:YFP* mice as a function of postnatal age. Animals were gavaged with tamoxifen at either 8 weeks, 30 weeks, or 1 year of age, then fed cuprizone for 6 weeks and perfused at 6 weeks recovery post cuprizone withdrawal. **F**, Schematic representation of a coronal section of the adult mouse brain illustrating positive (+) and negative (–) positions along the mediolateral axis relative to the dorsolateral corner of the V-SVZ, which we designated as position zero (0 µm) (relates to plots shown in G). **G**, Mean density of YFP^+^ CC1^+^ cells (NPC-derived oligodendrocytes) distributed along the mediolateral axis of the corpus callosum assessed at rostral, middle and caudal levels of the forebrain of *Nestin:YFP* mice as a function of postnatal age. Animals were gavaged with tamoxifen at either 8 weeks, 30 weeks, or 1 year of age, then fed cuprizone for 6 weeks and perfused at 6 weeks recovery post cuprizone withdrawal. Cell counts were performed for n=4-5 mice/group. Data represent mean ± SEM. Data were analyzed by two-way ANOVA with Tukey’s multiple comparisons test: \*\**p*<0.01, \*\*\*\**p*<0.0001. Scale bar: 200 µm (B). CC, corpus callosum; Ctx, cerebral cortex; LV, lateral ventricle; Sep, septum; Str, striatum.

To examine the effect of age on the production of NPC-derived oligodendrocytes following demyelination, we quantified the density of YFP^+^ CC1^+^ cells in the same sections. Cell counts revealed a prominent age-associated decline in the generation of NPC-derived oligodendrocytes (**Figure 2E**). Two-way ANOVA analyzing the effects of age and rostrocaudal position on YFP^+^ CC1^+^ cell density revealed a statistically significant interaction between the effects of age and rostrocaudal position (*F* (4, 33) = 10.78, *p*<0.0001). Simple main effects analysis showed that both age and rostrocaudal position had a statistically significant effect on YFP^+^ CC1^+^ cell density (*p*<0.0001).

Next, we plotted the local densities of YFP^+^ CC1^+^ cells along the mediolateral axis of the corpus callosum for each rostrocaudal position for mice of each age. We assigned the dorsolateral corner of the V-SVZ as the zero position (0 µm) and subdivided the corpus callosum into a series of 300 µm bins in both the medial (–‘ve) and lateral (+’ve) positions relative to this point (**Figure 2F**). We found the highest densities of YFP^+^ CC1^+^ cells in areas of the corpus callosum closest to the V-SVZ (**Figure 2G**). As the distance from the dorsolateral corner of the V-SVZ increased, the density of YFP^+^ CC1^+^ cells progressively decreased. This relationship was most obvious for cohorts aged 8 weeks or 30 weeks at experimental onset, although it also appeared to hold true for mice aged 1 year despite the overall minimal contribution of NPCs to oligodendrogenesis at this age.

### Intracisternal infusion of EGF or HB-EGF potentiated NPC-mediated oligodendrogenesis

Given the significant age-related decline in NPC contribution to oligodendrogenesis following cuprizone challenge, we theorized that the delivery of exogenous growth factors known to act on NPCs in the V-SVZ could potentiate NPC recruitment and oligodendrogenesis after cuprizone challenge. We elected to examine the effects of EGF, which acts via the EGF receptor, and heparin-binding EGF-like growth factor (HB-EGF), a member of the EGF family of growth factors that binds to and activates both EGFR and the related receptor tyrosine kinase, ErbB4. On the one hand, EGF delivery into the lateral ventricles has been demonstrated to promote NPCs in the V-SVZ to differentiate into OPCs and mature oligodendrocytes ^30^. Additionally, overexpression of human EGFR in CNP^+^ oligodendrocytes resulted in an increase in oligodendrocyte density and remyelination of the corpus callosum following lysolecithin-induced demyelination ^33^. On the other hand, infusion of HB-EGF into the lateral ventricles was shown to increase cell proliferation in the V-SVZ of aged mice ^22^. Whereas intranasal HB-EGF administration in young adult mice significantly increased cell proliferation in the V-SVZ and the mobilization of NPCs toward demyelinated lesions in the corpus callosum caused by lysolecithin injection ^29^.

To investigate whether exogenous delivery of EGF or HB-EGF can potentiate NPC recruitment and oligodendrogenesis following demyelination in the aged brain, we first examined the effects of growth factor infusion in mice aged 30 weeks at experimental onset. After 5 weeks of cuprizone challenge, *Nestin:YFP* mice received a 2-week intracisternal infusion of either EGF alone, HB-EGF alone, HB-EGF and EGF together, or aCSF (vehicle) (**Figure 3A**). The infusion was timed to overlap with the final week of cuprizone challenge and the first week of recovery following cuprizone withdrawal. We collected coronal brain sections from mice at 6 weeks recovery after cuprizone withdrawal and performed immunohistochemistry for YFP, SOX10 and CC1 together with a Hoechst nuclear counterstain. All growth factor-infused mice appeared to have higher densities of total YFP^+^ cells in the corpus callosum compared to vehicle-infused controls, particularly in the rostral and middle segments of the corpus callosum (**Figure 3B**). The increased density of NPC-derived cells was most obvious in mice that received a co-infusion of HB-EGF+EGF, and this was confirmed by counting YFP^+^ cell densities throughout the corpus callosum (**Figure 3C**). Two-way ANOVA analyzing the effects of infusion and rostrocaudal position on YFP^+^ cell density revealed a statistically significant interaction between the effects of infusion and rostrocaudal position (*F* (6, 84) = 13.77, *p*<0.0001). Simple main effects analysis showed that both age and rostrocaudal position had a statistically significant effect on YFP^+^ CC1^+^ cell density (*p*<0.0001). Tukey’s multiple comparisons tests revealed that all growth factor infusions significantly increased YFP^+^ cell densities in rostral segments whereas only co-infusion of HB-EGF+EGF significantly increased YFP^+^ cell densities in middle segments. Growth factor infusion did not significantly increase YFP^+^ cell density in the caudal segment of the corpus callosum.

**Figure 3.**
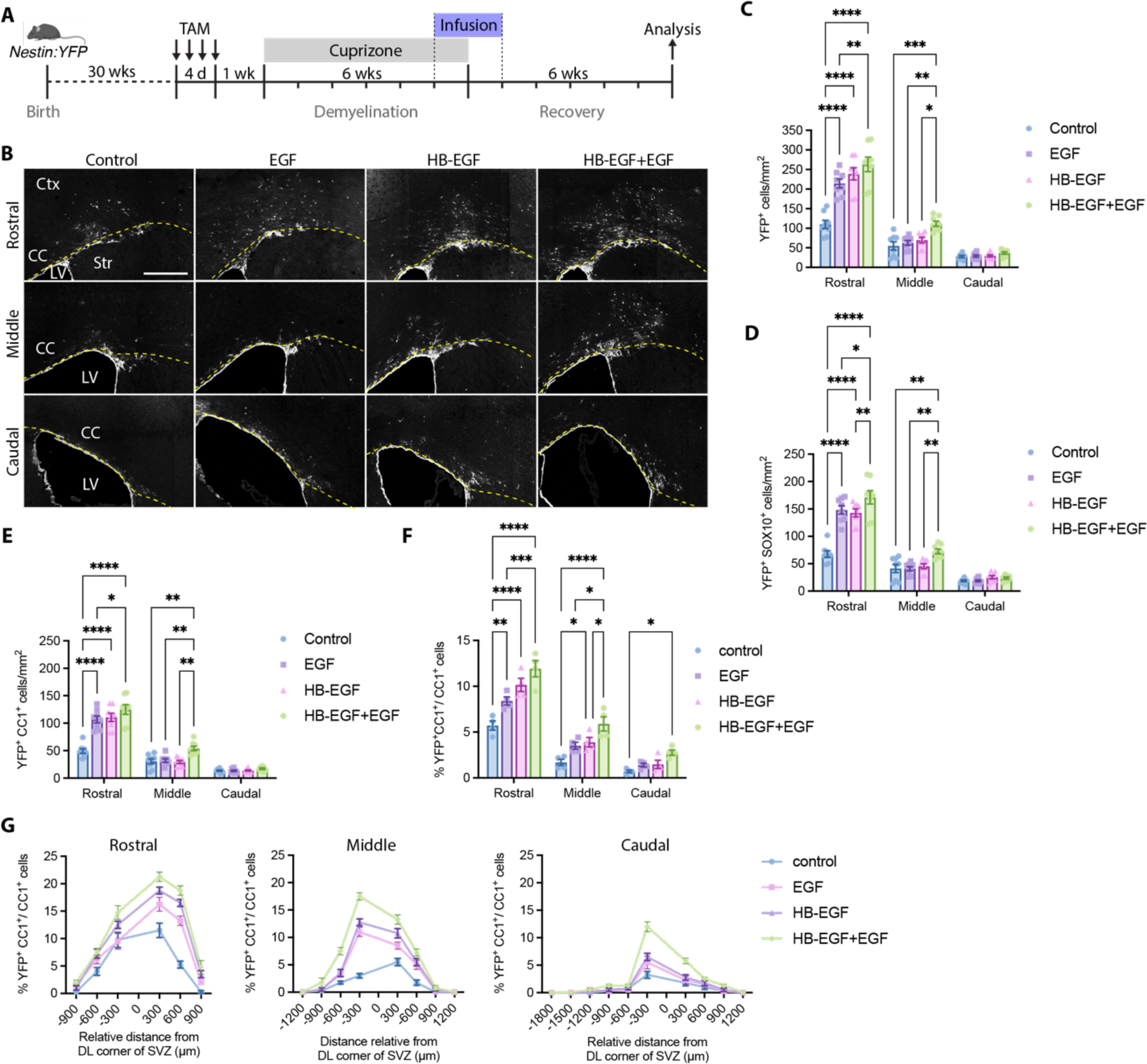
Intracisternal infusion of EGF or HB-EGF into the CSF of mice aged 30 weeks increased the density of NPC-derived cells in the corpus callosum assessed at 6 weeks recovery. **A**, Experimental timeline for *Nestin:YFP* mice indicating time points for tamoxifen gavage, cuprizone challenge, growth factors infusion and tissue collection. **B**, Confocal micrographs of coronal brain sections from 30-week-old *Nestin:YFP* mice assessed at 6 weeks recovery after cuprizone withdrawal. Images show YFP-expressing NPCs that have migrated from V-SVZ into the corpus callosum of mice infused with EGF, HB-EGF, HB-EGF+EGF, or vehicle control. The dashed yellow line indicates the ventral border of the corpus callosum. **C**, Density of total YFP^+^ cells in the corpus callosum at 6 weeks recovery. **D**, Density of YFP^+^ SOX10^+^ cells in the corpus callosum at 6 weeks recovery. **E**, Density of YFP^+^ CC1^+^ cells in the corpus callosum at 6 weeks recovery. **F**, The relative contribution of NPCs to oligodendrogenesis in rostral, middle and caudal segments of the corpus callosum. **G**, Plots of the % of CC1^+^ cells that co-express YFP according to their position along the mediolateral axis of the corpus callosum, where the dorsolateral corner of the V-SVZ is taken as 0 µm. Cell counts were performed for n=8 mice/group. Data represent mean ± SEM. Data were analyzed by two-way ANOVA with Tukey’s multiple comparisons tests: \**p*<0.05, \*\**p*<0.01, \*\*\**p*<0.001, \*\*\*\**p*<0.0001. Scale bar: 400 µm (B). CC, corpus callosum; Ctx, cerebral cortex; LV, lateral ventricle; Str, striatum.

Next, we quantified the densities of YFP^+^ SOX10^+^ cells in the corpus callosum of infused mice to determine the number of NPC-derived cells that had committed to the oligodendroglial lineage (**Figure 3D**). In the rostral segment, the density of YFP^+^ SOX10^+^ cells was more than two-fold higher in growth factor-infused mice compared to vehicle-infused controls (control: 67.6 ± 5.9 cells/mm^2^, EGF: 148.1 ± 7.7 cells/mm^2^, HB-EGF: 142.9 ± 7.1 cells/mm^2^, HB-EGF+EGF: 171.1 ± 12.0 cells/mm^2^, *p*<0.0001). In the middle segment, only the co-infusion with HB-EGF+EGF significantly increased the density of YFP^+^ SOX10^+^ cells (control: 41.2 ± 7.5 cells/mm^2^, HB-EGF+EGF: 72.5 ± 4.2 cells/mm^2^, *p*=0.0018). Growth factor infusion did not significantly alter the density of NPC-derived oligodendroglia in the caudal segment.

To establish whether growth factor infusion has an impact on the generation of NPC-derived oligodendrocytes following cuprizone challenge, we quantified the density of YFP^+^ CC1^+^ cells in the corpus callosum at 6 weeks recovery following cuprizone challenge in growth factor infused and control infused mice (**Figure 3E**). In the rostral segment, growth factor infusion significantly increased the density of YFP^+^ CC1^+^ cells compared to the controls (control: 49.3 ± 4.3 cells/mm^2^, EGF: 107.0 ± 6.3 cells/mm^2^, HB-EGF: 110.6 ± 7.5 cells/mm^2^, HB-EGF+EGF: 124.9 ± 8.8 cells/mm^2^, *p*<0.0001). In the middle segment, only the co-infusion with HB-EGF+EGF significantly increased YFP^+^ CC1^+^ cell density over control levels (control: 30.7 ± 4.5 cells/mm^2^, HB-EGF+EGF: 54.1 ± 4.0 cells/mm^2^, *p*=0.0028). Growth factor infusion did not significantly alter YFP^+^ CC1^+^ cell density in the caudal segment.

### Intracisternal infusion of EGF or HB-EGF increased the contribution of NPCs to oligodendrocyte regeneration after cuprizone challenge in 30-week-old mice

Given that growth factor infusion increased the production of NPC-derived oligodendrocytes in cuprizone-challenged mice aged 30 weeks, we asked whether this translated into an increase in the percentage contribution of NPCs-derived oligodendrocytes to the total oligodendrocyte population within the corpus callosum. We calculated the percentage of total CC1^+^ cells that expressed YFP, which represents the fraction of oligodendrocytes derived from fate-mapped NPCs (**Figure 3F**). YFP^−^ CC1^+^ cells in the corpus callosum 6 weeks after cuprizone withdrawal are either: 1) oligodendrocytes that survived the cuprizone challenge, 2) new-born oligodendrocytes that derive from pOPCs or 3) NPC-derived oligodendrocytes that escape genetic fate mapping, given that recombination in the *Nestin:YFP* line occurs with an efficiency of about ∼79% in mice aged 30 weeks. In the rostral segment of the corpus callosum, growth factor infusion significantly increased the contribution of NPCs to oligodendrocyte regeneration ∼1.5-fold in the EGF group and ∼2-fold for HB-EGF and HB-EGF+EGF groups (control: 5.71 ± 0.50%; EGF: 8.40 ± 0.45%; HB-EGF: 10.14 ± 0.72%; HB-EGF+EGF: 11.90 ± 0.89%; control vs EGF, *p*=0.0042; control vs HB-EGF or HB-EGF+EGF, *p*<0.0001). In the middle segment, both HB-EGF alone or HB-EGF+EGF co-infusion significantly increased NPC contribution (control: 1.69 ± 0.34%, HB-EGF: 3.89 ± 0.51%, HB-EGF+EGF: 5.89 ±0.78%; control vs HB-EGF, *p*=0.028; control vs HB-EGF+EGF, *p*<0.0001). In the caudal segment, HB-EGF+EGF infusion significantly increased NPC contribution in this segment (control: 0.70 ± 0.14%, HB-EGF+EGF: 2.76 ± 0.29%, *p*=0.038).

To understand whether growth factor infusion altered the distribution of NPC-derived oligodendrocytes across the mediolateral axis of the corpus callosum, we plotted the local percentage of YFP^+^ CC1^+^ cells relative to their distance from the dorsolateral corner of the V-SVZ, for rostral, middle and caudal segments (**Figure 3G**). This analysis revealed that the territories occupied by NPC-derived oligodendrocytes in growth factor-infused mice closely reflected those observed in vehicle-infused controls. We conclude that the increased density of NPC-derived oligodendrocytes in the corpus callosum of growth factor-infused mice is due to an increase in the recruitment of NPCs in the same areas that they are recruited in vehicle controls, rather than an expansion in their distribution across the mediolateral axis.

### Myelin density within the corpus callosum of Nestin:YFP mice aged 30 weeks was not changed by the infusion of growth factors during remyelination

To ascertain whether growth factor infusion following 5 weeks of cuprizone challenge influenced measures of myelin integrity within the corpus callosum at the 6-week recovery time point, we performed analyses to compare the abundance of myelin between groups. Compact myelin was visualized using spectral confocal reflectance (SCoRe) microscopy which clearly demarcated the corpus callosum and enabled the identification of bundled fiber tracts coursing across its mediolateral axis (**Figure 4A**). Quantitation of the mean optical density of SCoRe-labeled myelin within the corpus callosum was similar throughout its rostrocaudal axis and was not significantly altered by growth factor infusion (**Figure 4B**). We also examined myelin abundance by performing immunohistochemical detection of the two most abundant myelin proteins, proteolipid protein (PLP) and myelin basic protein (MBP) (**Figure 4C,E**). Optical density quantification of PLP and MBP expression in the corpus callosum revealed no significant differences in labeling intensity between treatment groups in the rostral, middle or caudal segments (**Figure 4D,F**). These data indicate that a two-week infusion of EGF and/or HB-EGF at the onset of remyelination does not measurably change myelin abundance when assessed at the 6-week recovery time point following cuprizone withdrawal.

**Figure 4.**
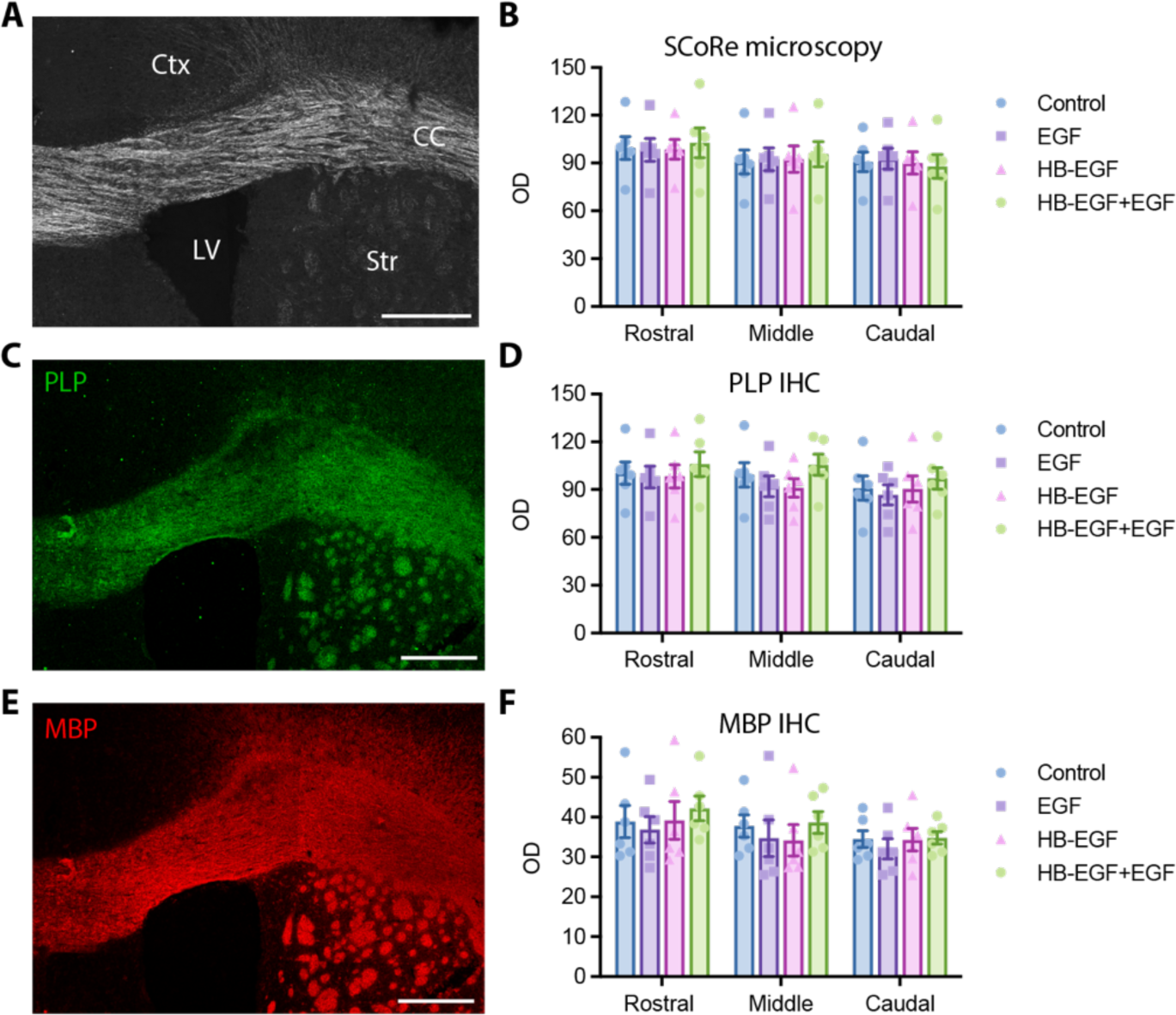
Growth factor infusions did not alter measures of myelin density in the corpus callosum of 30-week-old *Nestin:YFP* mice assessed at 6 weeks recovery after cuprizone withdrawal. **A,** Spectral confocal reflectance (SCoRe) microscopy of the rostral corpus callosum in an HB-EGF+EGF-infused mouse and optical density (OD). **B**, Quantification of SCoRe signal intensity from photomicrographs of the corpus callosum of growth factor-infused or vehicle-infused controls. Data are presented as mean optical density (OD) for rostral, middle and caudal segments. **C**, Confocal micrograph of a coronal brain section of an HB-EGF+EGF-infused mouse assessed at 6 weeks recovery following immunohistochemical detection of PLP. **D**, Plot of optical density measures of PLP labeling in the rostral, middle and caudal segments of the corpus callosum of growth factor-infused or vehicle-infused controls. **E**, Confocal micrograph of a coronal brain section of an HB-EGF+EGF-infused mouse assessed at 6 weeks recovery following immunohistochemical detection of MBP. **F**, Plot of optical density measures of MBP labeling in the rostral, middle and caudal segments of the corpus callosum of growth factor-infused or vehicle-infused controls. Data represent mean ± SEM. Scale bar: 300 µm (A,C,E). CC, corpus callosum; Ctx, cerebral cortex; LV, lateral ventricles; Str, striatum.

### Age reduced NPC recruitment to the corpus callosum after cuprizone challenge, but HB-EGF+EGF infusion at remyelination onset potentiated recruitment

Since the infusion of HB-EGF+EGF elicited the largest increase in the generation of NPC-derived oligodendrocytes in 30-week-old mice, we decided to explore the effects of HB-EGF+EGF infusion on mice of different postnatal ages. *Nestin:YFP* mice aged 8 weeks, 30 weeks, or 1 year were orally administered tamoxifen before being challenged with tamoxifen. Subsequently, they were infused with either HB-EGF+EGF or vehicle control for 2 weeks, starting on the last week of cuprizone challenge (**Figure 5A**). Animals were perfused after 6 weeks recovery following cuprizone withdrawal. Coronal brain sections were subjected to immunohistochemical processing to identify the presence of YFP, OLIG2, PDGFRA, CC1 and Hoechst (**Figure 5B**).

**Figure 5.**
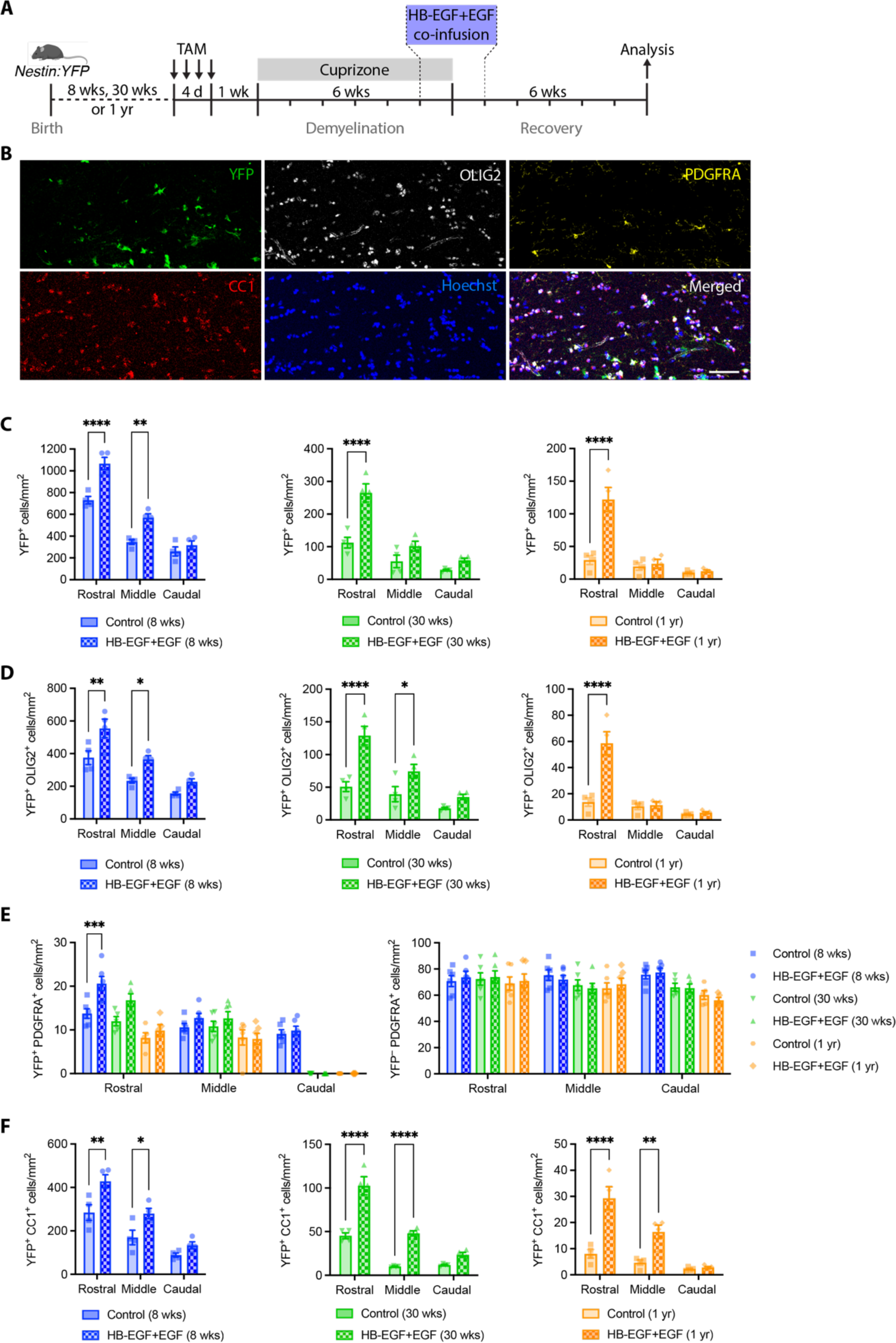
Infusion of HB-EGF+EGF into the CSF after cuprizone challenge attenuated the age-related decline in NPC-mediated oligodendrocyte regeneration. **A**, Experimental timeline for *Nestin:YFP* mice indicating time points for tamoxifen gavage, cuprizone challenge, growth factors infusion and tissue collection. **B**, Example confocal micrograph of an area of the corpus callosum of an HB-EGF+EGF-infused *Nestin:YFP* mouse taken from the 8-week cohort. Immunohistochemistry was performed against YFP, OLIG2, PDGFRA, CC1, and Hoechst to label NPC-derived cells, oligodendroglia, OPCs, oligodendrocytes, and cell nuclei, respectively. **C**, Density of YFP^+^ cells (NPC-derived cells) in the corpus callosum of *Nestin:YFP* mice aged 8 weeks, 30 weeks or 1 year that were infused with HB-EGF+EGF or vehicle control and assessed at 6 weeks recovery. **D**, Density of YFP^+^ OLIG2^+^ cells in the corpus callosum at 6 weeks recovery following cuprizone withdrawal shows the abundance of NPC-derived cells that have committed to the oligodendroglial lineage. **E**, Density of YFP^+^ PDGFRA^+^ cells (NPC-derived OPCs) and YFP^−^ PDGFRA^+^ cells (pOPCs) in the corpus callosum of mice at 6 weeks recovery. **F**, Density of YFP^+^ CC1^+^ cells in the corpus callosum at 6 weeks recovery after cuprizone withdrawal reveals the distribution of NPC-derived oligodendrocytes in the rostral, middle and caudal segments for each age group. Data represent mean ± SEM. Data were analyzed by two-way ANOVA with Tukey’s multiple comparisons tests: \**p*<0.05, \*\**p*<0.01, \*\*\**p*<0.001, \*\*\*\**p*<0.0001. Scale bar: 50 µm (B).

First, we quantified the density of NPC-derived YFP^+^ cells throughout the corpus callosum of vehicle-infused controls and HB-EGF+EGF-infused mice aged 8 weeks, 30 weeks, or 1 year (**Figure 5C**). Across all age groups, the infusion of HB-EGF+EGF increased the density of YFP^+^ cells in the rostral segment. In the 8-week cohort, the density of YFP^+^ cells increased significantly from 730.5 ± 33.5 cells/mm^2^ in the control group to 1066.2 ± 56.5 cells/mm^2^ in the HB-EGF+EGF-infused mice (*p*<0.0001). Similarly, for 30-week control vs 30-week HB-EGF+EGF animals, the density of YFP^+^ cells increased from 112.3 ± 16.6 cells/mm^2^ to 265.0 ± 27.9 cells/mm^2^ (*p*<0.0001). Finally, in 1-year control vs 1-year HB-EGF+EGF mice, the density of YFP^+^ cells increased from 29.2 ± 6.8 cells/mm^2^ to 122.1 ± 18.2 cells/mm^2^ (*p*<0.0001). In contrast, the infusion of HB-EGF+EGF only significantly increased the density of YFP^+^ cells in the middle segment for the 8-week cohort (8 wk control vs 8 wk HB-EGF+EGF, 344.7 ± 22.0 cells/mm^2^ vs 570.5 ± 35.9 cells/mm^2^, *p*=0.0022).

Next, we examined the effect of HB-EGF+EGF infusion on the generation of YFP^+^ OLIG2^+^ cells (NPC-derived oligodendroglial lineage cells) across different age groups (**Figure 5D**). The results showed a significant increase in the density of YFP^+^OLIG2^+^ cells in the rostral segment in all age groups (8-week control vs 8-week HB-EGF+EGF: 375.4 ± 42.4 cells/mm^2^ vs 555.0 ± 56.6 cells/mm^2^, *p*=0.0028; 30-week control vs 30-week HB-EGF+EGF: 51.0 ± 7.5 cells/mm^2^ vs 129.0 ± 14.1 cells/mm^2^, *p*<0.0001; 1-year control vs 1-year HB-EGF+EGF: 13.7 ± 3.2 cells/mm^2^ vs 58.6 ± 8.8 cells/mm^2^, *p*<0.0001). Similarly, in the middle segment, there was a significant increase in YFP^+^ OLIG2^+^ cell density in the 8-week and 30-week age groups (8-week control vs 8-week HB-EGF+EGF: 234.4 ± 15.0 cells/mm^2^ vs 365.1 ± 23.0 cells/mm^2^, *p*=0.0295; 30-week control vs 30-week HB-EGF+EGF: 39.4 ± 11.5 cells/mm^2^ vs 74.4 ± 10.7 cells/mm^2^, *p*=0.0480).

We also quantified the density of NPC-derived OPCs (YFP^+^ PDGFRA^+^ cells) and pOPCs (YFP^−^ PDGFRA^+^ cells) in the corpus callosum to assess whether HB-EGF+EGF infusion changed the balance between these two populations at the 6-week recovery time point after cuprizone withdrawal (**Figure 5E**). Whilst small numbers of YFP^+^ PDGFRA^+^ cells were observed in the rostral and middle segments at all ages, YFP^+^ PDGFRA^+^ cells were only observed in the caudal segment of animals aged 8 weeks. No NPC-derived OPCs were detected in the caudal segments of mice aged 30 weeks or 1 year, irrespective of the type of infusion.

We conducted a two-way ANOVA to compare the effect of infusion type and postnatal age on YFP^+^ PDGFRA^+^ cell density. The analysis revealed statistically significant effects of age on YFP^+^ PDGFRA^+^ cell density in each rostrocaudal segment (Rostral: *F* (2, 26) = 19.44, *p*<0.0001, Middle: *F* (2, 30) = 5.172, *p*=0.0118, Caudal: *F* (2, 30) = 192.3, *p*<0.0001) and infusion type on YFP^+^ PDGFRA^+^ cell density in the rostral segment (*F* (1, 26) = 16.43, *p*=0.0004). Post hoc analysis showed that compared to vehicle-infused controls, HB-EGF+EGF-infused mice aged 8 weeks had significantly higher densities of YFP^+^ PDGFRA^+^ cells in the rostral segment of the corpus callosum (8-week control vs 8-week HB-EGF+EGF, 13.7 ± 1.1 cells/mm^2^ vs 20.6 ± 1.6 cells/mm^2^, *p*=0.0004). We also analyzed the density of YFP^−^ PDGFRA^+^ cells (pOPCs) and found similar densities throughout the corpus callosum across groups. Two-way ANOVAs revealed no significant effect of infusion type or age on YFP^−^ PDGFRA^+^ cell density in rostral or middle segments. For the caudal segment, age had a statistically significant effect on the density of YFP^−^ PDGFRA^+^ cells (*F* (2, 26) = 18.73, *p*<0.0001).

Next, we examined the effect of HB-EGF+EGF infusion on the age-associated decline in the generation of NPC-derived oligodendrocytes. We plotted the density of YFP^+^ CC1^+^ cells in each segment of the corpus callosum for all age groups (**Figure 5F**). Two-way ANOVA revealed statistically significant effects of postnatal age (Rostral: *F* (2, 18) = 122.7, *p*<0.0001; Middle: *F* (2, 18) = 286.2, *p*<0.0001; Caudal: F (2, 18) = 282.2, *p*<0.0001) and infusion type (Rostral: *F* (1, 18) = 17.13, *p*=0.0006; Middle: *F* (1, 18) = 27.52, *p*<0.0001; Caudal: *F* (1, 18) = 18.62, *p*=0.0004) on the density of YFP^+^ CC1^+^ cells in the corpus callosum. There was also a statistically significant interaction between postnatal age and infusion type for the middle and caudal segments (Middle: *F* (2, 18) = 13.44, *p*=0.0003; Caudal: *F* (2, 18) = 9.395, *p*=0.0016). Sidak’s multiple comparisons test revealed the infusion of HB-EGF+EGF resulted in a significant increase in the density of YFP^+^ cells in the rostral and middle segments for each age group. The 8-week control group had a density of 284.4 ± 36.1 cells/mm^2^, while the 8-week HB-EGF+EGF group had a density of 428.2 ± 30.4 cells/mm^2^ (*p*=0.0038). Similarly, the 30-week control group had a density of 45.3 ± 3.4 cells/mm^2^, whereas the 30-week HB-EGF+EGF group had a density of 102.5 ± 10.4 cells/mm^2^ (*p*<0.0001). Moreover, the 1-year control group had a density of 8.0 ± 1.7 cells/mm^2^, whereas the 1-year HB-EGF+EGF group had a density of 29.3 ± 4.4 cells/mm^2^ (*p*<0.0001). In the middle segment, the density of YFP^+^ CC1^+^ cells was also significantly increased for each age group. The 8-week control group had a density of 169.8 ± 33.4 cells/mm^2^, while the 8-week HB-EGF+EGF group had a density of 279.9 ± 23.5 cells/mm^2^ (*p*=0.0270). Similarly, the 30-week control group had a density of 10.0 ± 0.4 cells/mm^2^, while the 30-week HB-EGF+EGF group had a density of 48.0 ± 2.9 cells/mm^2^ (*p*=0.0001). Additionally, the 1-year control group had a density of 4.7 ± 1.0 cells/mm^2^, while the 1-year HB-EGF+EGF group had a density of 16.4 ± 2.6 cells/mm^2^ (*p*=0.0032). Infusion did not result in a significant change in the caudal segment at any age.

### The density of total oligodendrocytes and myelin abundance in the corpus callosum of remyelinated mice was not altered by aging or HB-EGF+EGF infusion

To investigate the extent to which NPC-derived oligodendrocytes contribute to the overall regeneration of oligodendrocytes in the corpus callosum following cuprizone challenge, we first measured the density of total CC1^+^ cells in the corpus callosum for each age group, irrespective of YFP expression. Our findings showed that the infusion of HB-EGF+EGF did not significantly alter the density of total CC1^+^ cells in the corpus callosum for any age group (**Figure 6A**). We also measured the myelin abundance in the corpus callosum using SCoRe microscopy and immunohistochemistry against PLP and MBP, and found that it was similar across the rostrocaudal axis of the corpus callosum (**Figure 6B-D**). Two-way ANOVAs revealed that the type of infusion and postnatal age did not affect the density of CC1^+^ cells or the intensity of myelin in the corpus callosum for any age group, except for a statistically significant effect of postnatal age on MBP intensity specifically in the rostral segment (*F* (2, 30) = 6.002, *p*=0.0064). Based on these findings, we conclude that while HB-EGF+EGF infusion increased the abundance of NPC-derived mature oligodendrocytes in the corpus callosum, the total number of mature oligodendrocytes remained unchanged across different ages, and myelin abundance was similar in all groups.

**Figure 6.**
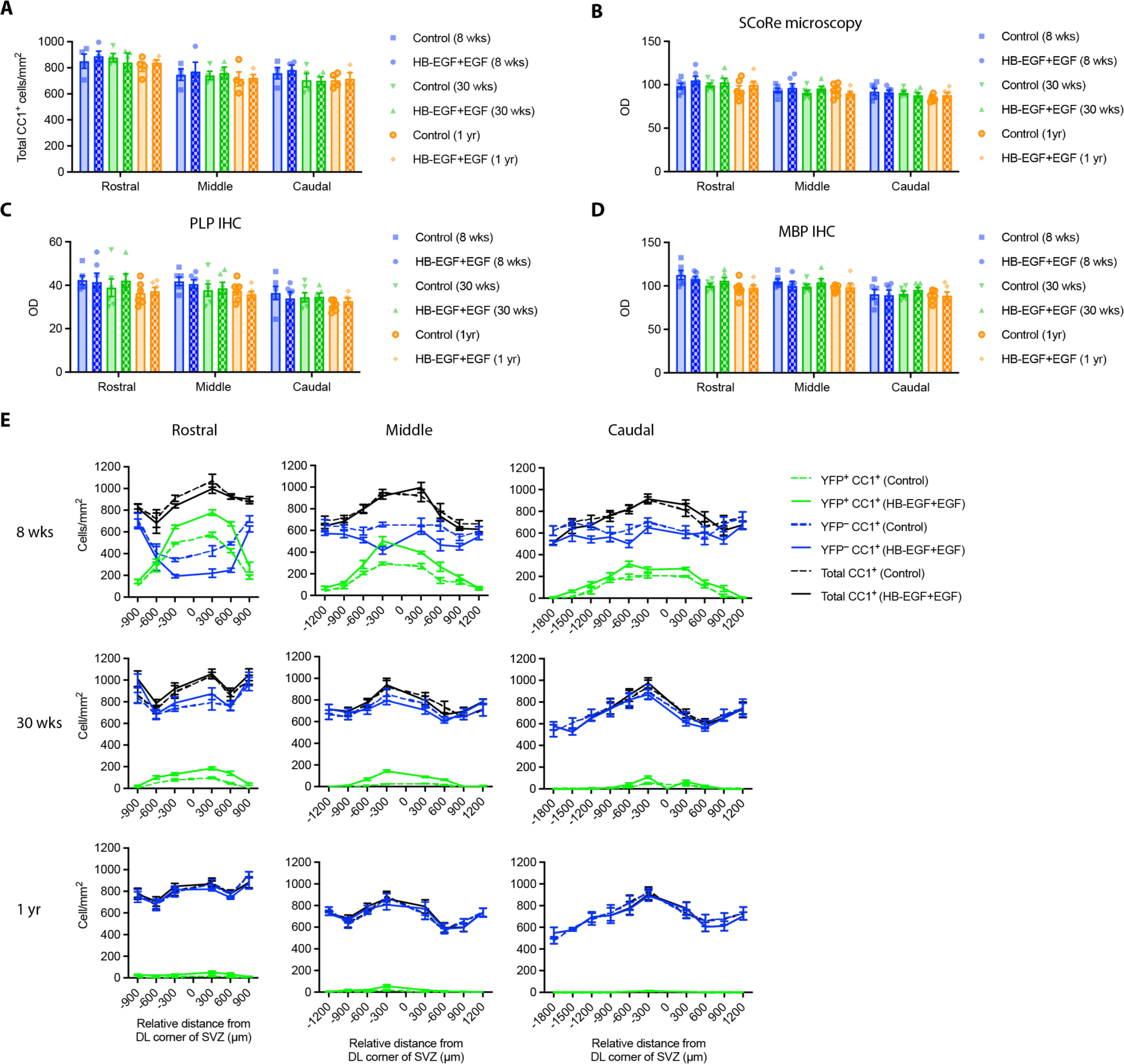
Postnatal age and HB-EGF+EGF infusion did not alter total oligodendrocyte density or measures of myelin abundance in the corpus callosum of *Nestin:YFP* mice assessed at 6 weeks recovery after cuprizone withdrawal. **A,** Density of total CC1^+^ oligodendrocytes (YFP^+^ and YFP^−^ fractions combined) in the corpus callosum of *Nestin:YFP* mice at 6 weeks recovery. **B**, Optical density of compact myelin in the corpus callosum measured from photomicrographs captured by spectral confocal reflectance microscopy. **C**,**D**, Optical density measures of PLP (C) and MBP (D) fluorescence intensity in the corpus callosum of acquired confocal images. **E,** Density of CC1^+^ oligodendrocytes across the mediolateral axis of the corpus callosum for the 8-week, 30-week and 1-year cohorts. The distributions of total CC1^+^ cells (black lines), YFP^+^ CC1^+^ cells (green lines) and YFP^−^ CC1^+^ cells (blue lines) are plotted separately for both HB-EGF+EGF-infused (solid lines) and vehicle-infused *Nestin:YFP* controls (dashed lines). Data represent mean ± SEM.

Next, we plotted the distribution of YFP^+^ CC1^+^ cells, YFP^−^ CC1^+^ cells, and total CC1^+^ cells across the mediolateral axis of the corpus callosum for each rostrocaudal segment and each age group to gain a better appreciation for the effects of NPC-mediated oligodendrogenesis on oligodendrocyte regeneration after cuprizone withdrawal (**Figure 6E**). Visualization of the cellular distributions demonstrated that the territories occupied by NPC-derived oligodendrocytes in HB-EGF-infused mice were very similar in pattern to that observed in vehicle-infused controls, albeit with exaggerated cellular recruitment. We also observed that in areas of greatest NPC recruitment, particularly in the rostral segment of HB-EGF+EGF-infused mice for the 8-week cohort, the uniform total density of CC1^+^ cells was achieved by a marked reduction in the density of YFP^−^ oligodendrocytes, likely reflecting a reduction in pOPC-mediated oligodendrogenesis in these areas. Interestingly, we also noted that the overall pattern of total oligodendrocyte distribution along the mediolateral axis appeared to be unique for each rostrocaudal segment and was largely maintained across each age group.

### HB-EGF+EGF infusion significantly increased the percentage contribution of NPC-derived oligodendrocytes to the total oligodendrocyte population

To determine the extent to which oligodendrocytes derived from neural progenitor cells (NPCs) contribute to the population of oligodendrocytes, we calculated the percentage of NPC-derived oligodendrocytes out of the total CC1+ cells in Nestin:YFP mice infused with HB-EGF+EGF or vehicle control, at each postnatal age and in each rostrocaudal segment (**Figure 7A**). We used two-way ANOVA to analyze the effects of age and infusion type on the % of CC1^+^ cells expressing YFP. The results showed that both age and infusion type had significant effects on the % contribution of YFP^+^ CC1^+^ cells to the total oligodendrocyte population. Specifically, HB-EGF+EGF infusion resulted in a significant increase in the % contribution of YFP^+^ CC1^+^ cells in all rostral segments at all age groups, and in all callosal segments for the 8-week cohort. Moreover, there was a statistically significant increase in the % contribution of YFP^+^ CC1^+^ cells in the middle segment for the 30-week cohort.

**Figure 7.**
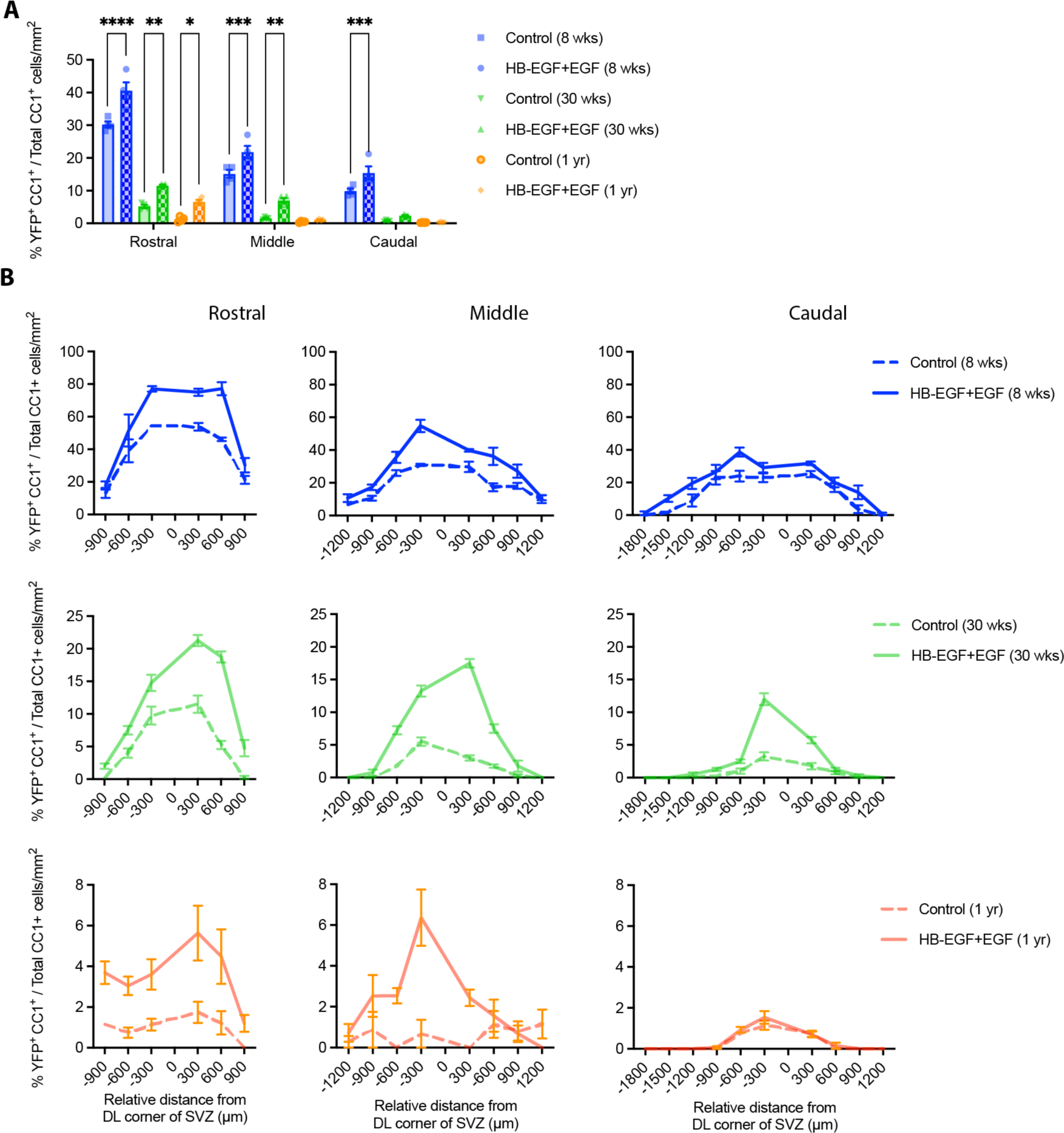
HB-EGF+EGF infusion significantly increased the contribution of NPC-derived oligodendrocytes to the total oligodendrocytes present in the corpus callosum at 6 weeks recovery after cuprizone challenge. **A**, Plot of the % of total CC1^+^ cells that express YFP, indicating the % contribution of NPC-derived oligodendrocytes to the total oligodendrocyte population in the corpus callosum of *Nestin:YFP* mice assessed at 6 weeks recovery following cuprizone withdrawal. **B**, Plots of the % of total CC1^+^ cells that express YFP across the mediolateral axis of the corpus callosum for the 8-week, 30-week and 1-year cohorts for HB-EGF+EGF-infused (solid lines) and vehicle-infused *Nestin:YFP* controls (dashed lines). Data represent mean ± SEM. Data (A) were analyzed by two-way ANOVA with Sidak’s multiple comparisons test: \**p*<0.05, \*\**p*<0.01, \*\*\*\**p*<0.0001.

Next, we assessed whether the infusion of growth factors affected the distribution of NPC-derived oligodendrocytes across the mediolateral axis of the corpus callosum. To analyze this, we measured the percentage of callosal CC1^+^ cells expressing YFP in relation to the distance from the dorsolateral corner of the V-SVZ (**Figure 7B**). Our data showed that the distribution of NPC-derived oligodendrocytes was similar between the growth factor-infused mice and the vehicle-infused controls. However, the regions that had the highest recruitment of NPCs in the vehicle-infused controls exhibited exaggerated contributions in the HB-EGF+EGF-infused mice. Therefore, we concluded that the increased density of NPC-derived oligodendrocytes in the corpus callosum of growth factor-infused mice is a result of a heightened response to demyelination, rather than an expansion in their distribution along the mediolateral axis.

### HB-EGF+EGF co-infusion did not alter the lineage specification of NPCs that migrate into the corpus callosum during remyelination

Our prior analyses focused on assessing the oligodendrogenic fate of NPC-derived cells in the corpus callosum. However, we have previously demonstrated that following cuprizone-induced demyelination, NPC-derived cells can become specified to alternate cell fates ^4^. To determine the fate of YFP^+^ cells, we performed immunohistochemistry using antibodies against four different markers: OLIG2, ALDH1L1, DCX, and PCNA (**Figure 8A**). The percentage of YFP^+^ cells that expressed OLIG2^+^, ALDH1L1^+^, and DCX^+^ was quantified to determine the extent of NPC differentiation into oligodendroglia, astrocytes, and neuroblasts, respectively (**Figure 8B**). Across all age groups and treatment conditions, approximately 40-65% of migrated NPCs differentiated into oligodendroglia, 25-40% differentiated into astrocytes, and 1-4% differentiated into neuroblasts which are predominately observed in the caudal segment. Additionally, between 4-17% of YFP^+^ cells in the corpus callosum remained unidentified, that is, they did not express OLIG2, ALDH1L1, or DCX.

**Figure 8.**
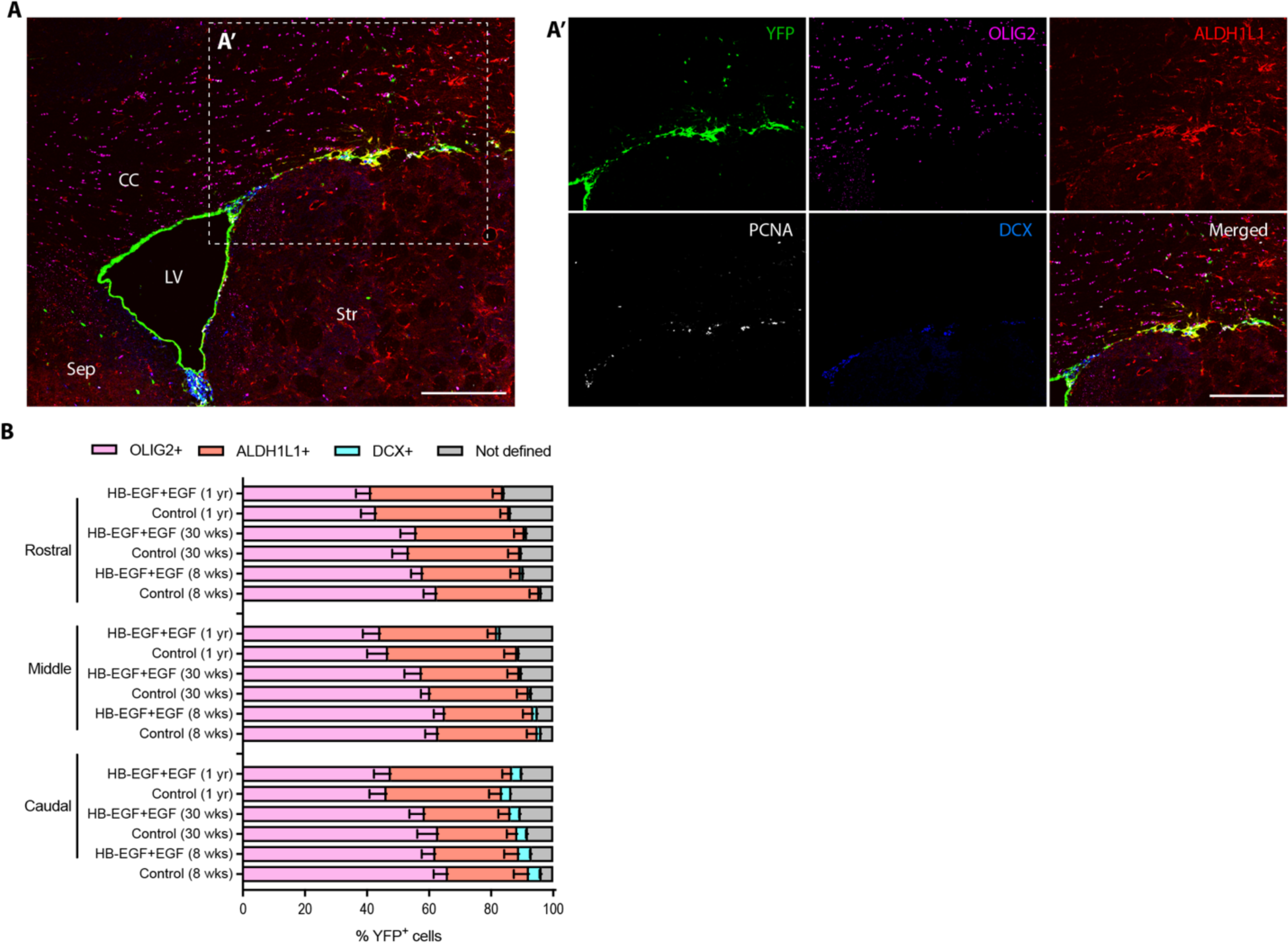
Aging influenced the cell fate specification of NPC-derived cells during remyelination of the corpus callosum but lineage specification was not altered by infusion with HB-EGF+EGF. **A**, Representative confocal micrograph of a coronal brain section processed immunohistochemically to detect YFP, ALDH1L1, OLIG2, DCX and PCNA to label NPC-derived cells, astrocytes, oligodendroglia, neuroblasts and dividing cells, respectively. **A’**, High magnification of the dashed box shown in A. **B**, The % of NPC-derived cells in the corpus callosum of *Nestin:YFP* mice at 6 weeks recovery that expressed oligodendroglial (OLIG2^+^), astroglial (ALDH1L1^+^) or neuronal (DCX^+^) markers. YFP^+^ cells that did not express any of these markers were classified as ‘not defined’. Data represent mean ± SEM. Scale bar: 300 µm (A, A’). CC, corpus callosum; LV, lateral ventricle; Sep, septum; Str, striatum.

Two-way ANOVA on data for the % YFP^+^ cells that expressed OLIG2^+^ revealed no statistically significant effects of rostrocaudal position or infusion, but a statistically significant effect of postnatal age (Rostral: *F* (2, 24) = 5.746, *p*=0.0091; Middle: *F* (2, 24) = 7.434, *p*=0.0031; Caudal: *F* (2, 24) = 7.628, *p*=0.0027). Similarly, two-way ANOVA on data for the % YFP^+^ cells that expressed ALDH1L1^+^ revealed no statistically significant effects of infusion, but a statistically significant effect of postnatal age for rostral and caudal segments (Rostral: *F* (2, 30) = 4.865, *p*=0.0148; Caudal: *F* (2, 30) = 3.692, *p*=0.0369).

Finally, two-way ANOVA on data for the % YFP^+^ cells that expressed DCX^+^ revealed no statistically significant effects of age or infusion type, but a statistically significant effect of rostrocaudal position (8 week: *F* (2, 24) = 32.41, *p*<0.0001; 30 week: *F* (2, 42) = 15.05, *p*<0.0001; 1 year: *F* (2, 24) = 38.42, *p*<0.0001). These data suggest that the specification of NPCs to an oligodendroglial fate declines with postnatal age and is not altered by HB-EGF+EGF infusion.

### The number of dividing NPCs in the V-SVZ decreased significantly with advancing age, but increased following HB-EGF+EGF infusion

To investigate the extent to which the proliferation in the V-SVZ is influenced by postnatal age and growth factor infusion, we quantified the total number of PCNA^+^ cells in the dorsolateral corner of the V-SVZ of one hemisphere using the same brains used to examine NPC fate within the corpus callosum. Cell counts revealed a marked reduction in PCNA^+^ cells in the V-SVZ in sections collected at middle and caudal segments (**Figure 9A**). Two-way ANOVAs to test the effect of postnatal age and growth factor infusion revealed a statistically significant effect of postnatal age and infusion on PCNA^+^ cell number on the V-SVZ in both rostral segments (Age: *F* (2, 30) = 27.34, *p*<0.0001; Infusion type: *F* (1, 30) = 17.46, *p*=0.0002) and middle segments (Age: *F* (2, 30) = 47.23, *p*<0.0001; Infusion type: *F* (1, 30) = 20.72, *p*<0.0001; Interaction: *F* (2, 30) = 5.046, *p*=0.0129). Further, Sidak’s *post hoc* analysis revealed that HB-EGF infusion significantly increased PCNA^+^ cell numbers in 8-week-old mice in both rostral and middle segments (Rostral: control vs HB-EGF+EGF, 36.5 ± 3.1 cells vs 51.5 ± 6.1, *p*=0.0107; Middle: 9.8 ± 0.8 vs 17.2 ± 1.7, *p*<0.0001), and in 30-week old mice in the rostral segment (Rostral: control vs HB-EGF+EGF, 20.7 ± 1.1 vs 34.7 ± 3.5, *p*=0.0182).

**Figure 9.**
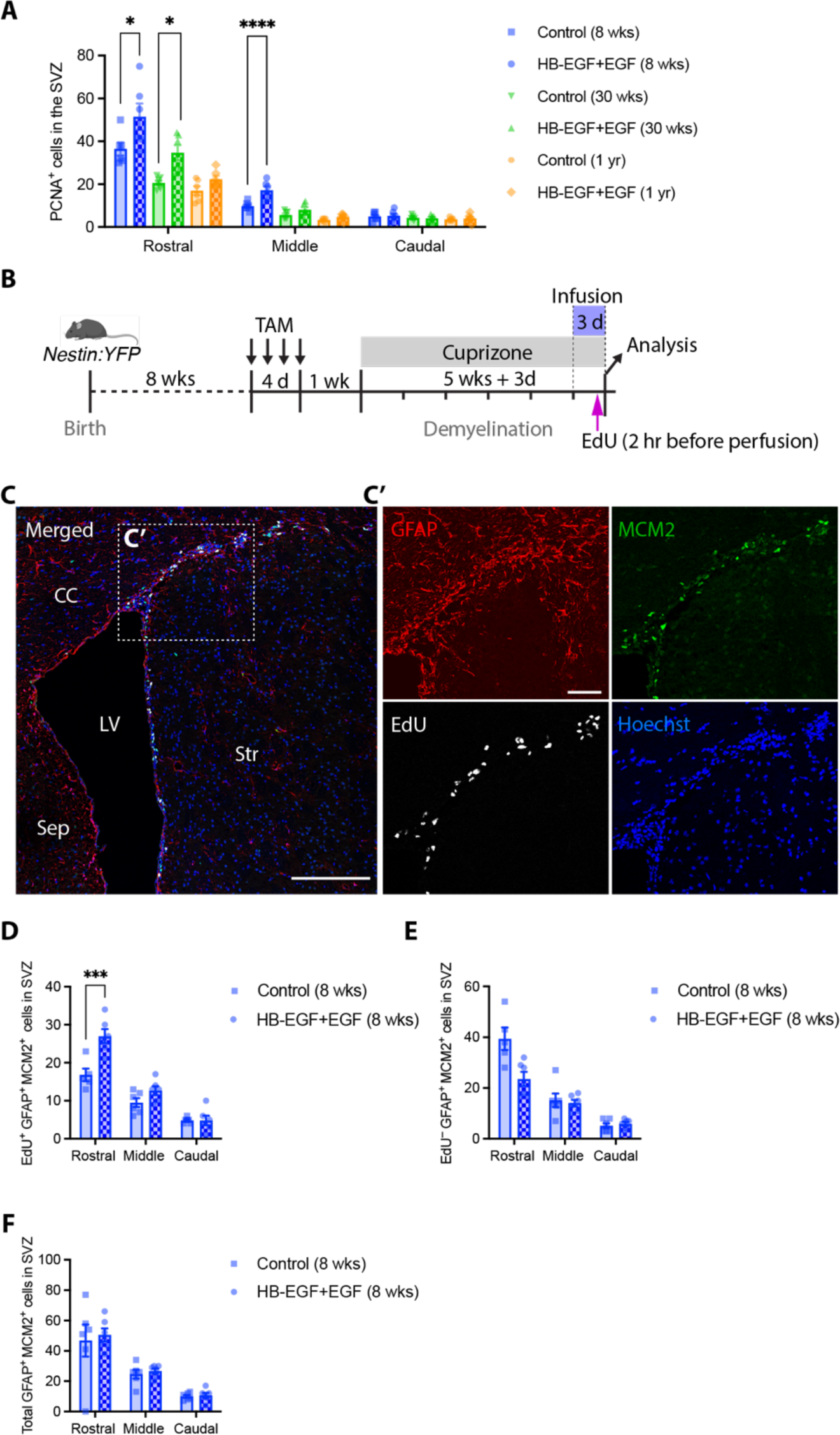
HB-EGF+EGF infusion increases the proliferation of NPCs and GFAP^+^ MCM2^+^ neural stem cells in the V-SVZ. **A**, Density of mitotic NPCs (PCNA^+^ cells) in the V-SVZ of *Nestin:YFP* mice at 6 weeks recovery after cuprizone withdrawal assessed across the rostocaudal axis for all age groups. **B**, Experimental timeline for *Nestin:YFP* mice indicating time points for tamoxifen gavage, cuprizone challenge, growth factor infusion, EdU injection and tissue collection. **C**, Representative confocal micrograph of a coronal brain section following immunohistochemistry to detect GFAP, MCM2, EdU and Hoechst. Labeling was performed to identify putative neural stem cells (NSCs) (GFAP^+^ MCM2^+^ cells) that passed through the S phase of the cell cycle within 2 hours after EdU injection. **C**, Absolute number of EDU^+^ GFAP^+^ MCM2^+^ cells (dividing NSCs) in the dorsolateral of the V-SVZ in rostral, middle and caudal segments. **D**, Absolute number of EDU^−^ GFAP^+^ MCM2^+^ cells (non-dividing NSCs) in the dorsolateral of the V-SVZ. Data represent mean ± SEM. Data were analyzed by two-way ANOVA with Sidak’s multiple comparisons test: \**p*<0.05, \*\*\**p*<0.001, \*\*\*\**p*<0.0001. Scale bar: 200 µm (C), 50 µm (C’). CC, corpus callosum; LV, lateral ventricle; Sep, septum; Str, striatum.

### Rapidly dividing NPCs in the V-SVZ were responsive to HB-EGF+EGF infusion

To investigate whether HB-EGF+EGF infusion increases the proliferation of neural stem cells in the V-SVZ, 8-week-old mice were given tamoxifen and then fed cuprizone for 5 weeks. After that, mice received an infusion of either vehicle control or HB-EGF+EGF for 3 days before the brains were harvested (**Figure 9B**). Two hours before perfusion/fixation, the mice received an intraperitoneal injection of EdU to label cells in the S phase of the cell cycle. Immunohistochemistry was performed against GFAP, MCM2, and Hoechst to label NPCs, mitotic cells, and cell nuclei, respectively (**Figure 9C**). EdU^+^ cells were detected using a Click-IT reaction. First, we quantified the number of EdU^+^ GFAP^+^ MCM2^+^ cells localized in the dorsolateral corner of the V-SVZ. In the rostral segment, HB-EGF infusion resulted in a statistically significant increase in the number of EdU^+^ GFAP^+^ MCM2^+^ cells in the dorsolateral corner of the V-SVZ (control vs HB-EGF+EGF, 16.8 ± 1.7 cells vs 27.0 ± 1.9 cells, p<0.0001) (**Figure 9D**). No significant differences were observed in other segments. We also quantified the number of EdU^−^ GFAP^+^ MCM2^+^ cells in the dorsolateral corner of the V-SVZ, which represent putative neural stem cells (NSCs) that did not enter the S phase during the period of EdU exposure (**Figure 9E**). Two-way ANOVA revealed statistically significant effects of both infusion type and rostrocaudal position on the number of EdU^−^ GFAP^+^ MCM2^+^ cells in the V-SVZ (Rostrocaudal position: *F* (2, 31) = 41.86, *p*<0.0001; Infusion: *F* (1, 31) = 5.094, *p*=0.0312). Next, we plotted the total number of GFAP^+^ MCM2^+^ cells present in the dorsolateral corner of the V-SVZ (**Figure 9E**). Interestingly, the total number of presumptive NSCs remained unaffected by growth factor infusion. Overall, these data support the conclusion that the infusion of HB-EGF+EGF led to the proliferation of putative NSCs in the V-SVZ.

## Discussion

Multiple Sclerosis (MS) is a common neurodegenerative disease, where immune cells infiltrate the CNS and destroy the myelin sheaths surrounding the axons ^1^. This leads to axonal degeneration and neurological dysfunction. Following demyelination, remyelination occurs, where pOPCs or NPCs migrate into the demyelinated lesions, generate new oligodendrocytes, and produce new myelin sheaths around demyelinated axons ^3–5^. Recent studies have shown that NPCs residing in the V-SVZ have the potential to generate new oligodendrocytes during the remyelination process ^4, 5, 34–36^. However, remyelination failure is common among MS patients ^37^, and the rate of remyelination declines significantly with aging ^9^. The effects of aging on the neurogenic capacity of NPCs are well studied, but the changes in the proliferation of these cells over time remain controversial ^14–18^, and it remains unclear how NPC-derived oligodendrogenesis during remyelination is influenced by age.

In this study, we investigated the effects of aging on NPC behavior following demyelination, as well as the effects of EGF and/or HB-EGF infusion on NPCs in different age groups. The V-SVZ niche plays a crucial role in regulating NPCs behavior ^19^, and changes in the expression level of signaling molecules and growth factors, including EGF, may contribute to the decline in NPC potential in the aged brain ^20–25^. Previous studies have shown the potential of EGF or HB-EGF in manipulating NPCs to contribute more to neurogenesis ^22^ or oligodendrogenesis ^29, 30, 33^, but most of these studies used young animal models (8-week-old mice), and their findings may not be applicable to older animals. Therefore, we chose EGF and HB-EGF to investigate their potential for manipulating NPCs following demyelination and restoring the contribution of these cells in an aged brain.

First, we examined the effects of intracisternal infusion of EGF, HB-EGF, and HB-EGF+EGF on the behavior of NPCs in 30-week-old *Nestin:YFP* mice that were challenged with cuprizone to induce CNS demyelination. Our data demonstrated that these treatments significantly increased the contribution of NPCs to oligodendrogenesis assessed 6 weeks following demyelination. The most effective treatment on 30-week-old mice was co-infusion of HB-EGF+EGF, so we used this treatment for 8-week-old and 1-year-old mice to examine the interaction between aging and the effects of HB-EGF+EGF infusion. We compared the behavior of NPCs in 8-weeks, 30-weeks, and 1-year-old *Nestin:YFP* mice, six weeks after cuprizone-induced demyelination. Our findings revealed that the proliferation of NPCs and their density in the corpus callosum declined dramatically with aging, indicating that the relative contribution of these cells to oligodendrogenesis declined significantly in older animals compared to younger ones. Moreover, no NPC-derived OPCs were observed in the caudal segment of the corpus callosum in either 30-week-old or 1-year-old mice. We demonstrated that infusion of HB-EGF and EGF into the CSF following demyelination increased the proliferation of NPCs in the V-SVZ and their contribution to oligodendrogenesis across all age groups. The rostral segment of the corpus callosum showed more pronounced improvement compared to the middle and caudal segments. Although the growth factors enhanced the contribution of NPCs to oligodendrogenesis, they had no influence on the total number of oligodendrocytes or the total amount of myelin in the corpus callosum at the 6-week recovery time point. These observations suggest that oligodendrocyte density in the corpus callosum is under tight homeostatic control. In regions where we observe increased recruitment of YFP^+^ NPC-derived oligodendrocytes there was a concomitant reduction in the density of YFP^−^ oligodendrocytes, supporting the notion that oligodendrogenic NPCs and pOPCs compete for remyelination of demyelinated axons in the corpus callosum ^38^.

Another key finding of this study was that oligodendroglial lineage specification of NPCs declined with age whereas astrogliogenic fate increased. Some studies have suggested that EGF potentiates the differentiation of V-SVZ-derived NPCs into oligodendrocytes ^30^. Interestingly, our results do not support a role for these factors in altering the lineage specification of NPCs, at least in the context of CNS demyelination. Rather, our findings support the possibility that co-infusion of HB-EGF+EGF increases the proliferation of neural stem cells and their progeny in the V-SVZ, thereby increasing the number of oligodendrogenic NPCs available to remyelinate the corpus callosum. Our findings align with a previous study that also showed the effect of these growth factors on the reactivation of neural stem cell quiescence in aged mice ^22, 39^.

Collectively, our findings reveal the significant impact of aging on the behavior of NPCs during the remyelination process. The data highlight the beneficial effects of growth factor infusion in enhancing the contribution of NPCs to oligodendrogenesis and indicate the therapeutic potential of HB-EGF and EGF to combat the negative effects of aging on remyelination efficacy.

## Acknowledgments

We acknowledge the Monash Micro Imaging (MMI) and the Monash Ramaciotti Centre for Cryo-Electron Microscopy. KM is the recipient of a Melbourne International Research Scholarship (MIRS). TDM was supported by a Future Fellowship from the Australian Research Council (FT150100207, T.D.M.), generous philanthropic support provided by Metal Manufactures Ltd., and was supported (in part) by the Intramural Research Program of the NIMH (Annual report: ZIAMH002985). The Australian Regenerative Medicine Institute is supported by grants from the State Government of Victoria and the Australian Government.

